# CREID – A ChemoReceptor-Effector Interaction Database

**DOI:** 10.1101/2023.05.04.539426

**Authors:** Vincent Peta, Timothy Hartman, Shiva Aryal, Bichar Shrestha Gurung, Ram Singh, Samuel Hass, Alain Bomgni, Tuyen Do, Saurabh Sudha Dhiman, Venkataramana Gadhamshetty, Etienne Z. Gnimpieba

## Abstract

The ChemoReceptor-Effector Interaction Database (CREID) is a collection of bacterial chemoreceptor and effector protein and interaction data to understand the process that chemoreceptors and effectors play in various environments. Our website includes terms associated with chemosensory pathways to educate users and those involved in collaborative research to help them understand this complex biological network. It includes 2,440 proteins involved in chemoreceptor and effector systems from 7 different bacterial families with 1,996 chemoeffector interactions. It is available at https://react-creid.bicbioeng.org.

**Key Highlights:**

1. CREID links bacterial chemoreceptors with their associated effectors.
2. Researchers interested in what attracts or repels bacteria can use CREID as a comprehensive source for information.
3. Biosensor developers can leverage CREID to discover better interactions for their applications.
4. CREID reveals knowledge gaps in chemoreceptor-effector interactions for both model and non-model organisms.

## Chemoreceptors & Effectors in the Environment

Bacterial chemosensory systems are critical in the movement behaviors of bacterial species, known as chemotaxis. Bacterial chemotaxis, movement under the influence of a chemical gradient, either towards (positive chemotaxis) or away (negative chemotaxis) to find optimum conditions for growth and survival due to chemoattractants or chemorepellents respectively. These chemosensory-signaling pathways allow the bacteria to sense the environment around them and travel toward or away from areas of stimuli, such as carbon sources and toxic chemicals. Chemotaxis could be based on bacterial behavioral responses such as energy taxis (stimuli affecting cellular energy), phototaxis (light) (Taylor & Zhulin, 1998), aerotaxis (oxygen concentration in niche) (Taylor, 1983), redox taxis (Armitage, 1997) and taxis to alternative electron acceptors (Grishanin et al., 1997).

The same stimulant could be attractant and repellent for two different bacteria or change in concentration could also reverse its effect from attractant to repellent (Shioi et al., 1987). Chemoreceptors are multi-domain proteins that act as input-signal receivers (that bind to ligands) in the environment and transfer it inside the cell to start the mechanism of response such as chemotaxis (Figure 1). Different stimuli (chemoeffectors) elicit unique responses which results in bacterial movement towards a specific stimulus (Bi & Lai, 2015a; Keegstra et al., 2022).

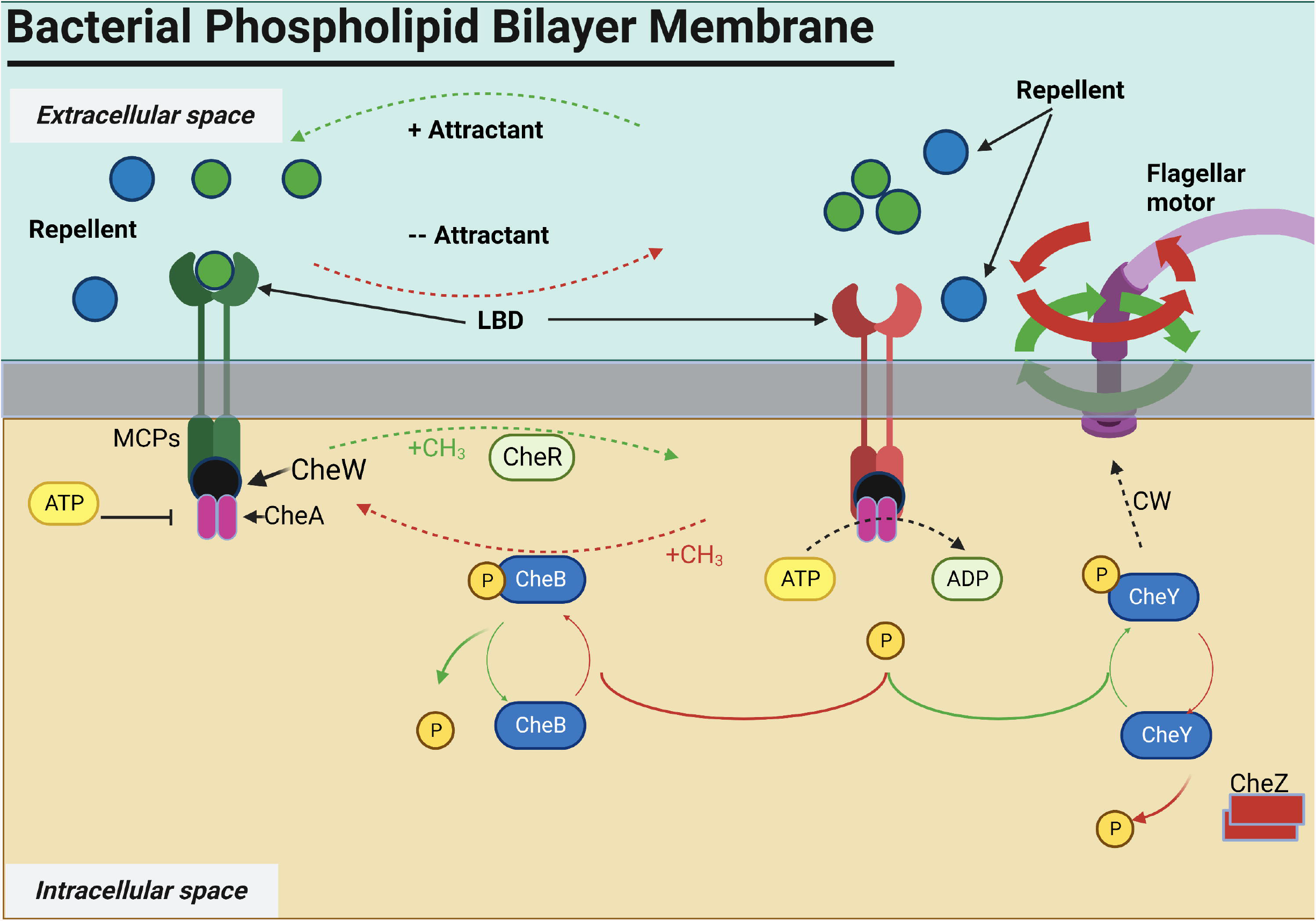

In marine environments multiple communities such as phytoplankton, zooplankton, and viruses co-exist with microbes (photoautotrophs, photoheterotrophs and heterotrophs) (Figure 2.). These communities have interactions at multiple levels for nutritional exchange. Marine environments are dynamic due to multiple factors such as nutrients which vary depending on the water current, sunlight, upwelling of sea water to various other factors (pH, temperature, salinity, oxygen, pressure) (Brumley et al., 2020). At the same time, nutrient dynamics can also be influenced by the microbial population themselves, which can augment the dissolved organic material (DOM) and inorganic substrates in the water (Blackburn et al., 1997; Jackson, 1980; McCarthy & Goldman, 1979; Smriga et al., 2016). These microenvironments (phycosphere (100 um) and the diffusion boundary layer are rich in significant amounts of simple sugars, amino acids, organic acids, complex polysaccharides, and lipids and are active zones for heterotrophic bacteria (Aaronson, 1978; Fogg, 1977; Hellebust, 1965; Jones & Cannon, 1986; Sharp, 1977). For example, *Synechococcus* sp. CS-94 RRIMP N1 releases 34 nitrogen containing compounds in marine environments and *Marinobacter adhaerens* HP15 metabolize 27 compounds (out of 34) released from strain CS-94 RRIMP N1 (Raina et al., 2023).

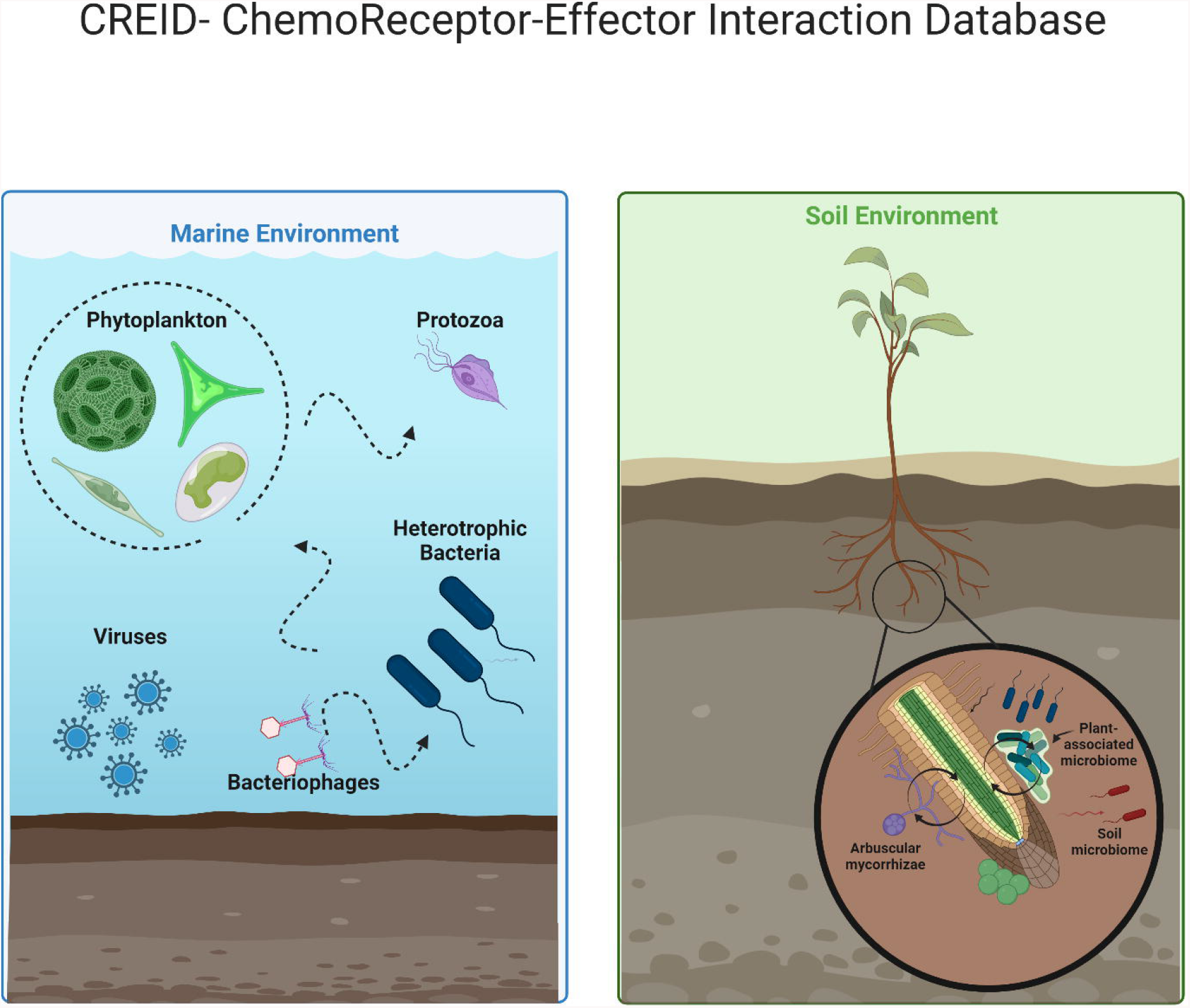

In soil systems, plant-microbe interaction is very critical. Plants recruit microbes by releasing root-exudates for microbial nutritional-supply and defense against pathogens. The chemical composition of root exudates has a direct effect on the rhizosphere communities nearest to the plant-host. These include plant growth-promoting rhizobacteria (PGPR) genera such as *Bacillus, Pseudomonas, Enterobacter, Burkholderia, Arthrobacter*, and *Paenibacillus* (Sasse et al., 2017; Zhang et al., 2017). For example, the banana root exudate fumaric acid attracts the Gram-positive *Bacillus subtilis* N11 and stimulates biofilm formation (Zhang et al., 2014); malic acid exuded by *Arabidopsis* stimulates binding to roots and biofilm formation on roots by *B. subtilis* strain FB17 (Rudrappa et al., 2008). Bacterial growth and antifungal activity of certain species of Gram-negative *Pseudomonas* spp. is dependent on organic acids and sugars isolated from tomato root exudates (Kravchenko et al., 2003; Mhlongo et al., 2018). Among PGPR, species of *Pseudomonas* and *Bacillus* are the best studied model organisms for beneficial plant-microbe interaction (Weller et al., 2002; Raaijmakers et al., 2010).

Beneficial bacteria use chemotaxis to move toward plant root chemoeffectors (exudates) by means of chemoreceptor pathways. The plant pathogen *Agrobacterium fabrum* uses chemotaxis to infect dicotyledon plants which includes roses and geraniums (Wang et al., 2021). Wang et al discovered the chemoreceptor protein responsible for this chemotaxis. With these findings the community can leverage this discovery to create new antimicrobial agents for dicotyledons or seeds that have two embryonic leaves. As more microbiologists unveil new chemoreceptor-effector interactions, more microbe-attracting and repelling agents can be discovered—even those that target specific genes that code for chemoreceptors.

## Environmental Factors and Conditions

Chemoeffectors occur both naturally and synthetically in an environmental system as any molecule or energy that either attracts or repels microbes (Bi & Lai, 2015b). This includes inorganic elements (metal ions), compounds (organic, aromatic), light, oxygen level and heat. These chemoeffectors could behave differently with variation in micro-niche conditions such as pH, salinity, humidity, barometric pressure, dissolved organic matter, distance between chemoeffectors and microbes, water turbulence, and depth of aquatic environment. Chemosensory pathways drive not just chemotaxis but also biofilm formation in a crosstalk fashion. Mediation of biofilm formation on can be influenced by various conditions in solid, fluid, and aerial environments, which can impact how bacterial species congregate and interact in biofilms (Boyeldieu et al., 2020; Huang et al., 2019). It is well established that chemotaxis play an important role in biofilm formation(O’Toole & Kolter, 1998; Pratt & Kolter, 1998; Prigent-Combaret et al., 1999; Watnick & Kolter, 1999) guiding bacterium to swim toward nutrients (chemoeffectors/chemoattractants) adsorbed to a surface followed by flagellar attachment to initiate the biofilm formation (Stelmack et al., 1999). By linking chemoreceptor proteins to environmental chemoeffectors, researchers can test environmental changes to modify microbe behavior in-turn biofilm formation and track proteomic and genomic changes that could lead to future studies.

## Biosensor Selection Criteria

Biosensors are devices or probes that integrates a biological element with an electronic component to generate a measurable signal. Naresh and Lee outlined six key characteristics of biosensors (Naresh & Lee, 2021). Table 1 explains these characteristics and how CREID can assist biosensor developers.

**Table 1.** Biosensor characteristics and how CREID assists biosensor developers match them.

|  |  |  |
| --- | --- | --- |
| Characteristic | Definition | How CREID Assists |
| *Selectivity* | Important for selecting a chemoreceptor or effector that is integrated into the biosensor. Receptors/effectors attached to a biosensor can detect the analyte of interest in a sample, even if the sample has impurities. | Researchers can link bioreceptors (chemoreceptors) of interest to analytes (chemoeffectors) and vice versa. |
| *Sensitivity* | “The minimum amount of analyte that can be correctly detected/identified in a minimum number of steps and in low concentrations (ng/mL or fg/mL) to verify the existence of analyte traces in the sample.” | Researchers can find relevant information to synthesize analytes more easily and quickly. CREID can also be used for computational chemistry and retrosynthesis via access to more available data without the need for manual searching. |
| *Linearity* | “Contributes to the accuracy of the measured results. The higher the linearity (straight line), the higher the substrate concentration detection.” | Populating CREID with new receptor and effector discoveries can improve accuracy through finding the most reliable interaction sets. |
| *Response Time* | The time needed to obtain a majority of the targeted result. | Users can leverage CREID in their experimental design to consider ideal environmental conditions for their protein reactions of choice. |
| *Reproducibility* | “Characterized by precision (similar output when the sample is measured more than once) and accuracy (capability of a sensor to generate a mean value closer to the actual value when the sample is measured every time). It is the ability of the biosensor to produce identical results whenever the same sample is measured more than once.” | Populating CREID with new receptor and effector discoveries can improve accuracy and precision through finding the most reliable interaction pairs/sets. |
| *Stability* | Important when continuous monitoring is required. Stability is how vulnerable the biosensor and its components are to environmental disturbances. Stability can be affected by the binding energy of the chemoreceptor to a chemoeffector, and receptor/effector degradation over the biosensor’s life. | Data and text mining in CREID can find the best substrates and materials to bind receptors and effectors in place without being washed away by other compounds. Users may find chemoreceptors, effectors, and substrates with the highest molecular binding energies to maximize stability and biosensor life. |
| Source: Naresh & Lee, 2021 |  |  |

## Data Selection

Data of chemosensory proteins as retrieved from databases: PubMed, Genbank (Benson et al., 2005), Uniprot (UniProt, 2019), Entrez (Maglott et al., 2011) and PROSITE (de Castro et al., 2006). Manual-based search extraction and curation were performed to fully account for chemosensory related proteins which could have previously been submitted as chemoeffector protein, not defined as a chemoattractant or chemorepellent. Confirmation steps were also used to remove any duplicates or low confidence/ predicted protein data. Retrieved data was transformed to structured data that can be searchable and downloaded in a CSV format. Data will also be available for download via API and other web applications.

## Database Organization

The focus of the CREID database (Figure 3.) is at the protein level that can be searched by microbial taxa, interaction type, and accession ID for easy data parsing and extraction. The database contains URLs for protein information in Uniprot, ligand binding domain information in the RCSB protein database, chemical information in PubChem, and the article containing the chemoreceptor-effector interaction discovery (Figure 4.)(Figure 5.). Uniprot was also used to confirm protein domains of suspected chemosensory associated proteins. Network visualization option can connect specific receptors to the STRING (Szklarczyk et al., 2019) to show protein-protein interaction network map and connect specific effectors to the STITCH (Kuhn M et al., 2008) to show protein-chemical interaction network map.

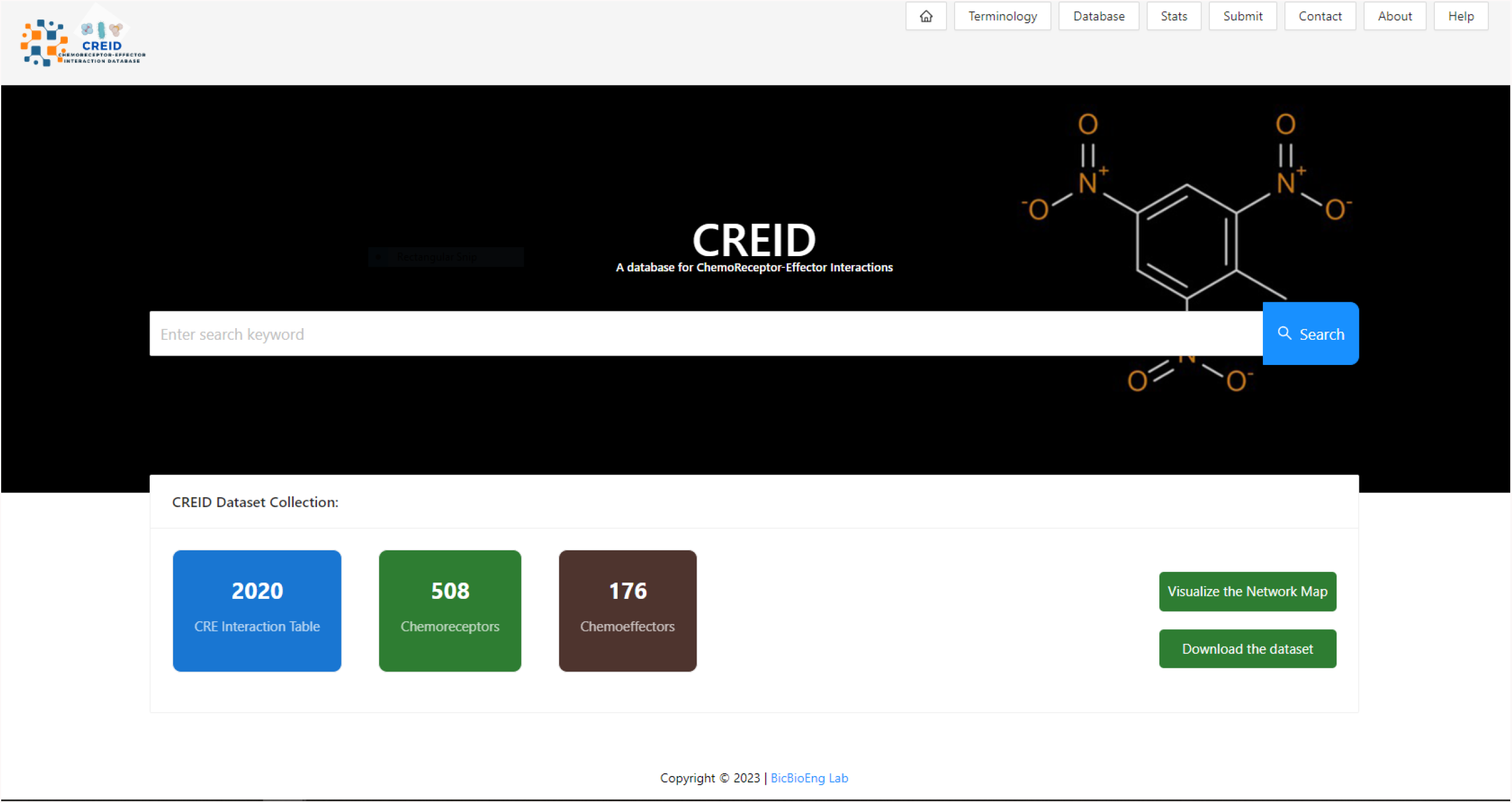

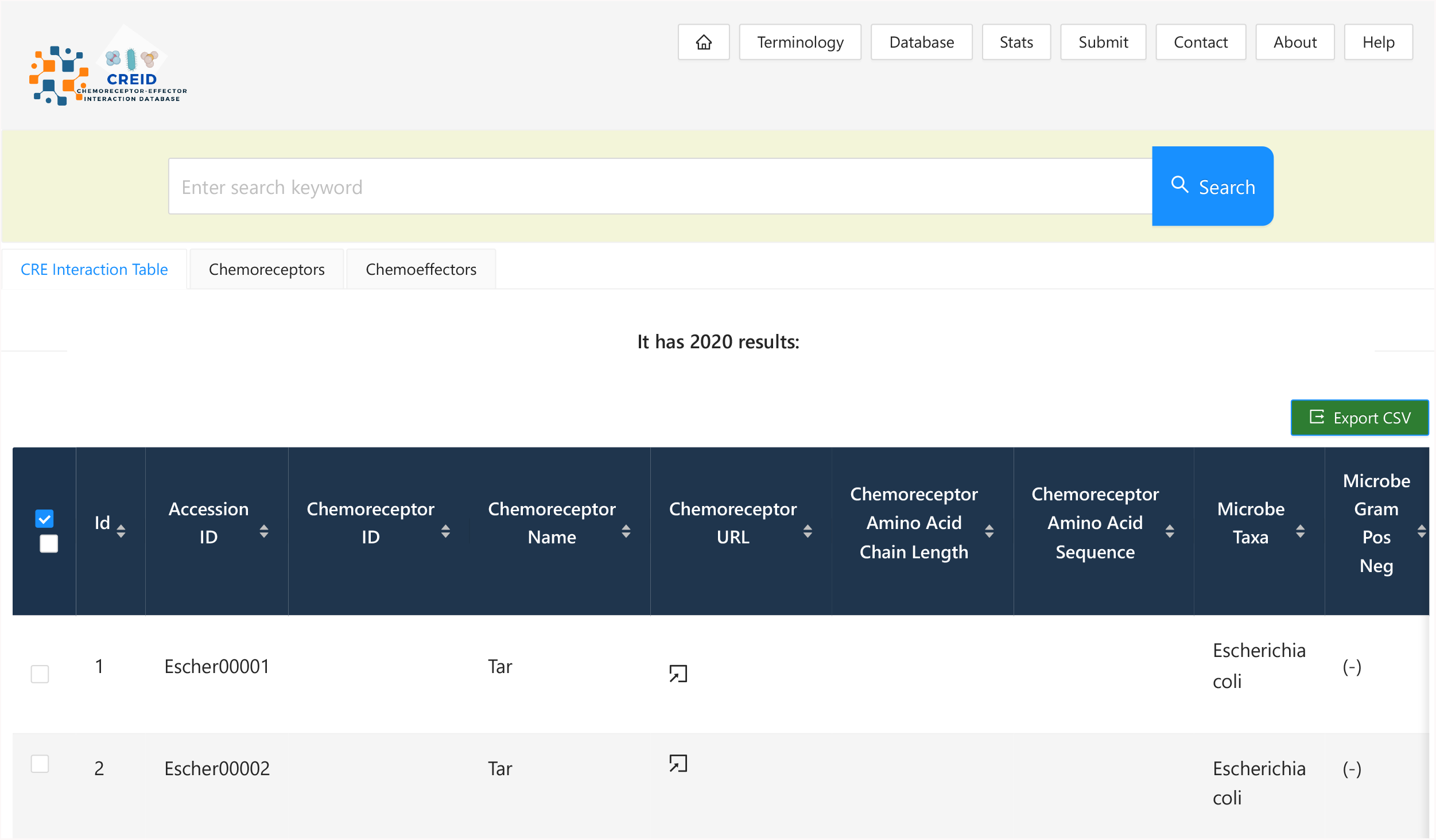

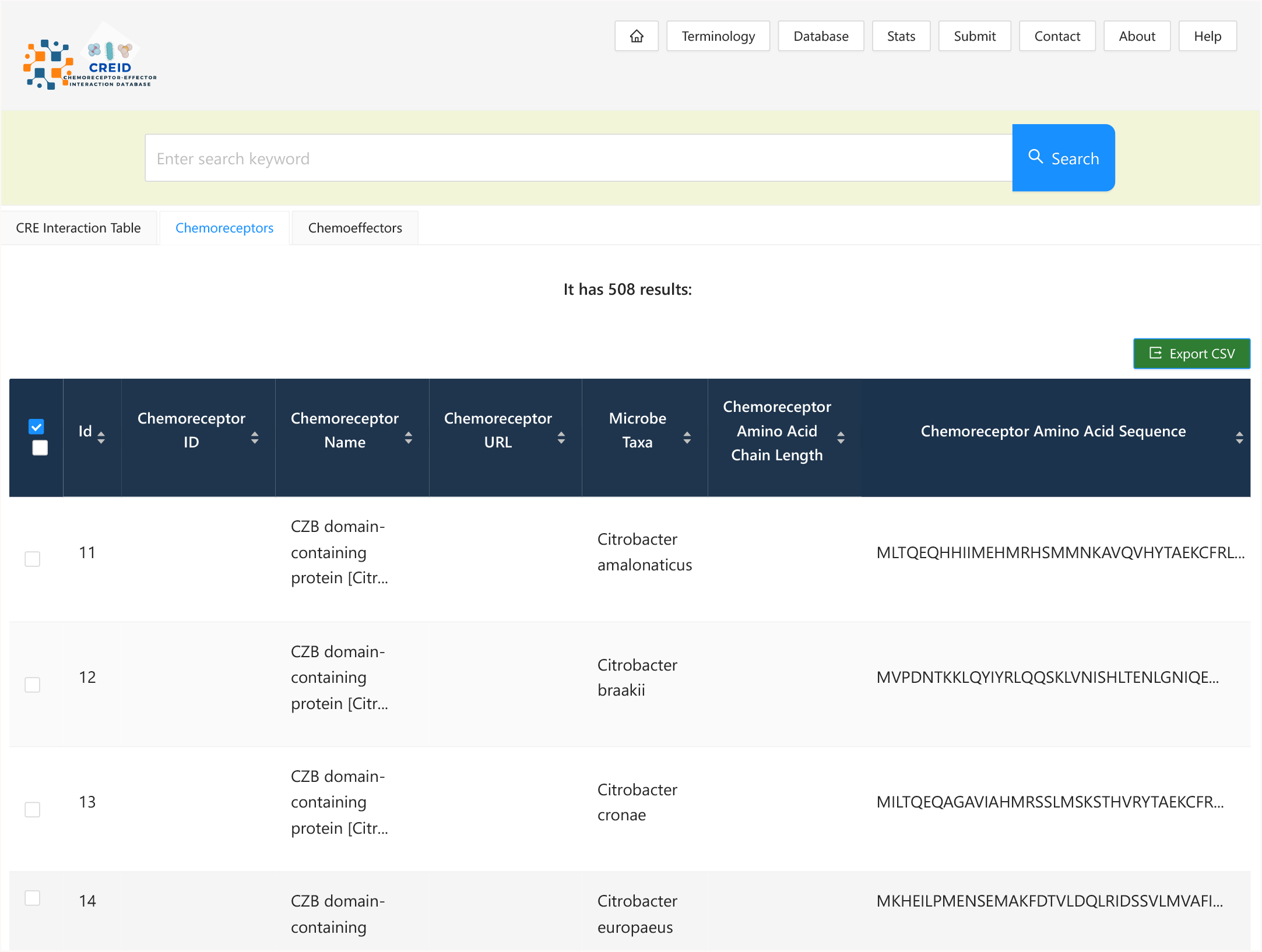
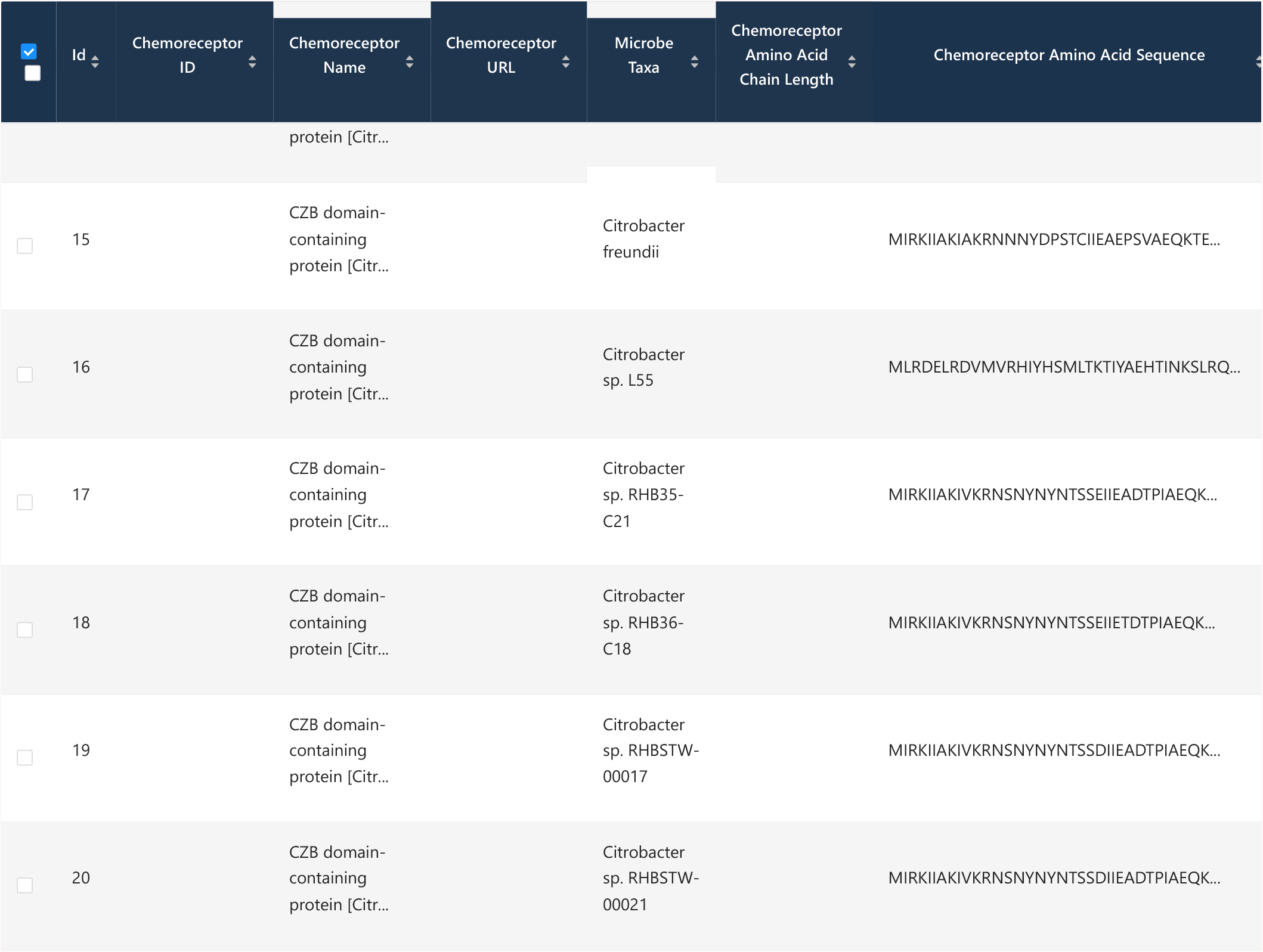
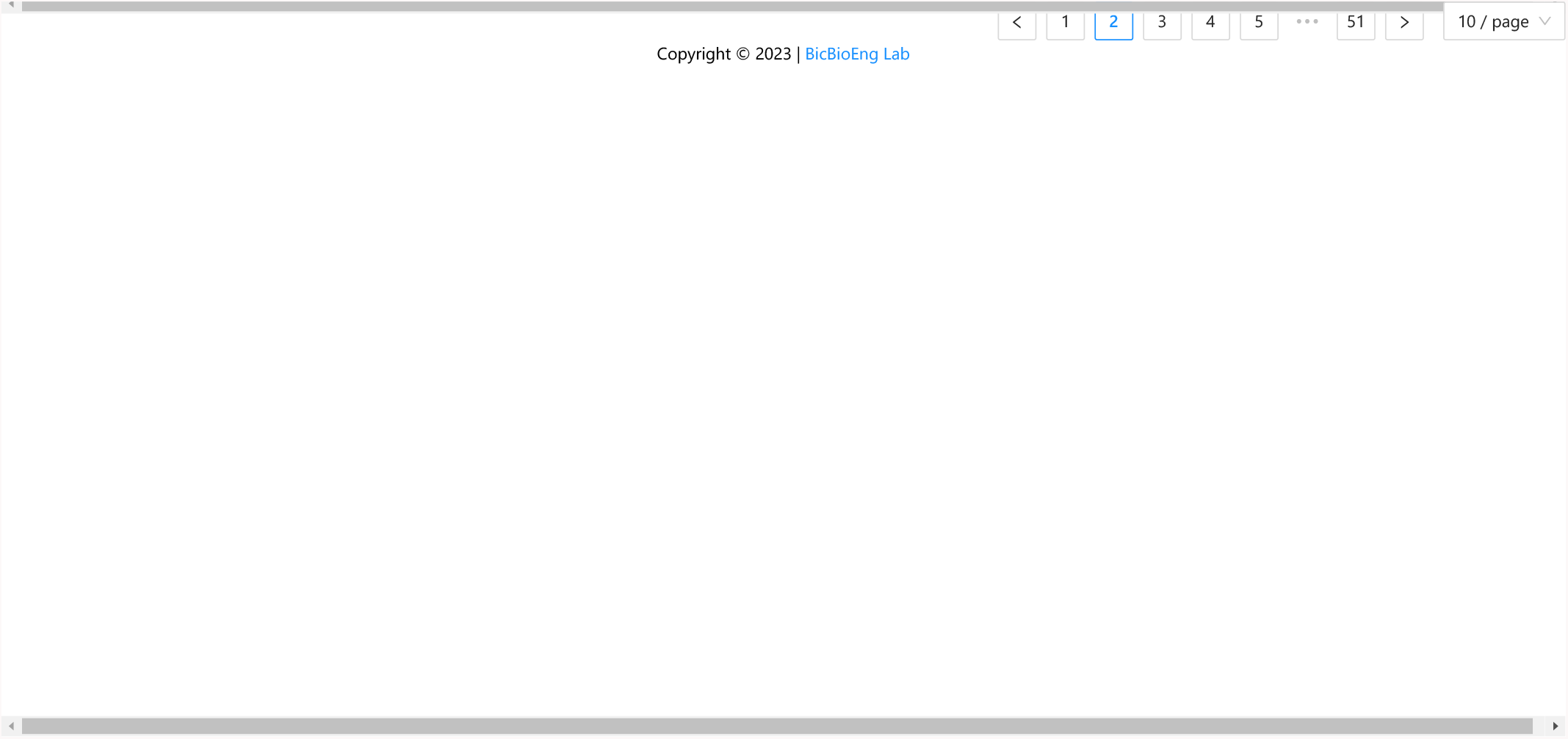

## Concluding Remarks

CREID is a free webserver and catalogue that contains known protein data for chemosensory proteins that are associated with chemotaxis of various bacterial taxa. It was designed to be used as a tool to collect, organize, and aid in understanding the complexities of chemosensory pathway interactions in different environments that anyone could use and target for future research.

## Supporting information

Supplemental GIF

## Acknowledgements & Funding

Research reported in this publication was supported by an Institutional Development Award (IDeA) from the National Institute of General Medical Sciences of the National Institutes of Health under grant number P20GM103443. The content is solely the responsibility of the authors and does not necessarily represent the official views of the National Institutes of Health. VP and TH are both first co-authors of this manuscript.

---

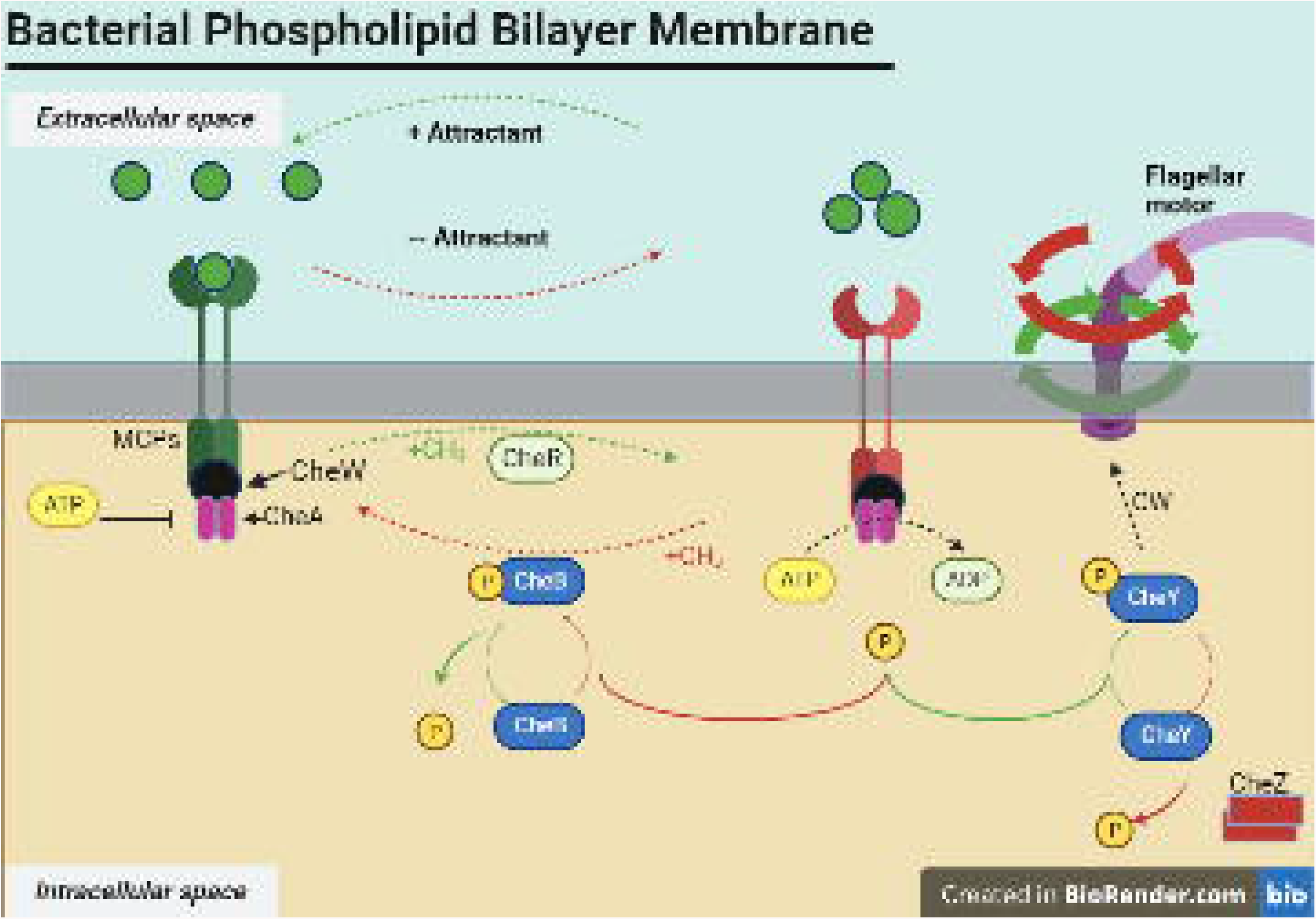

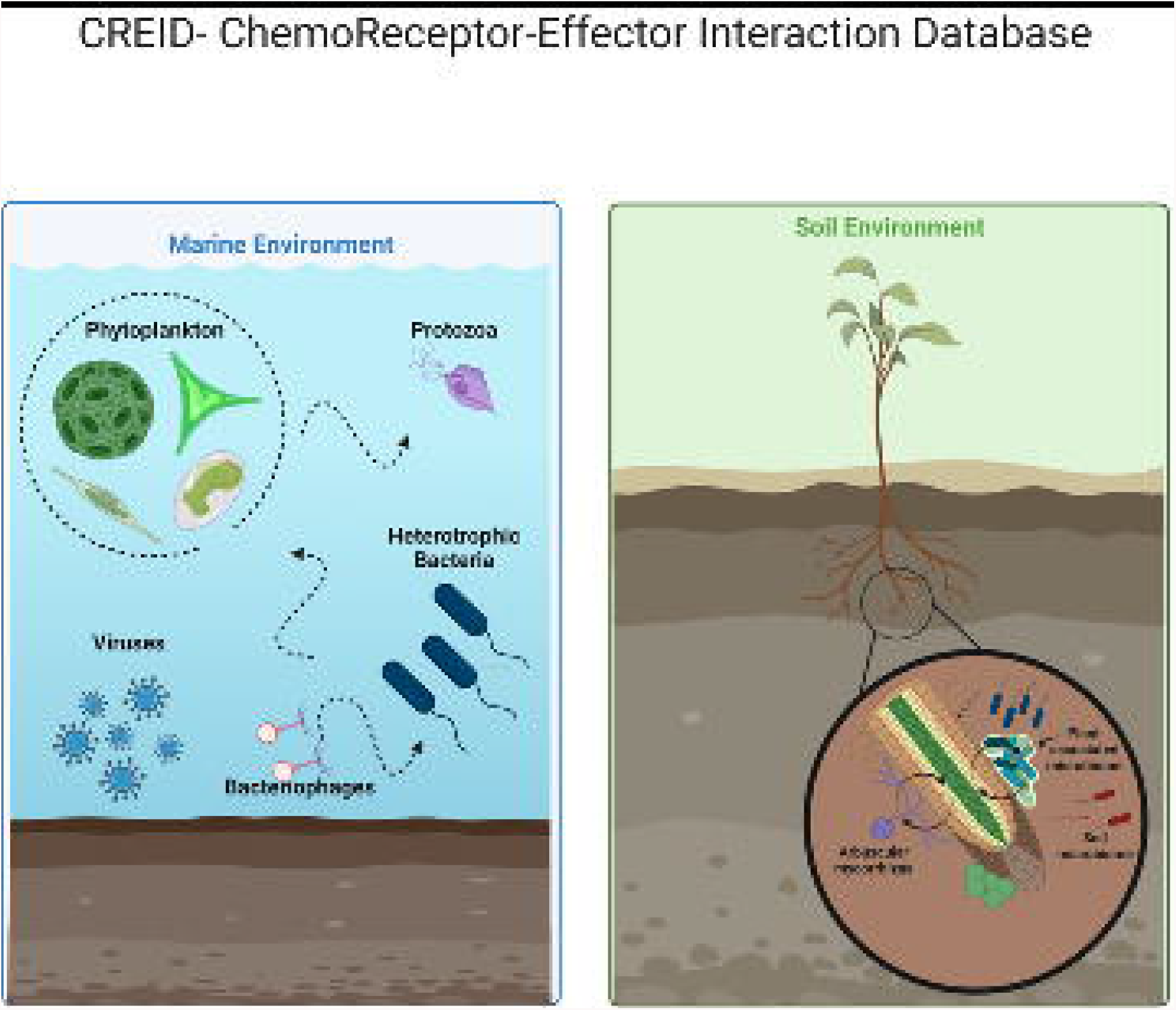

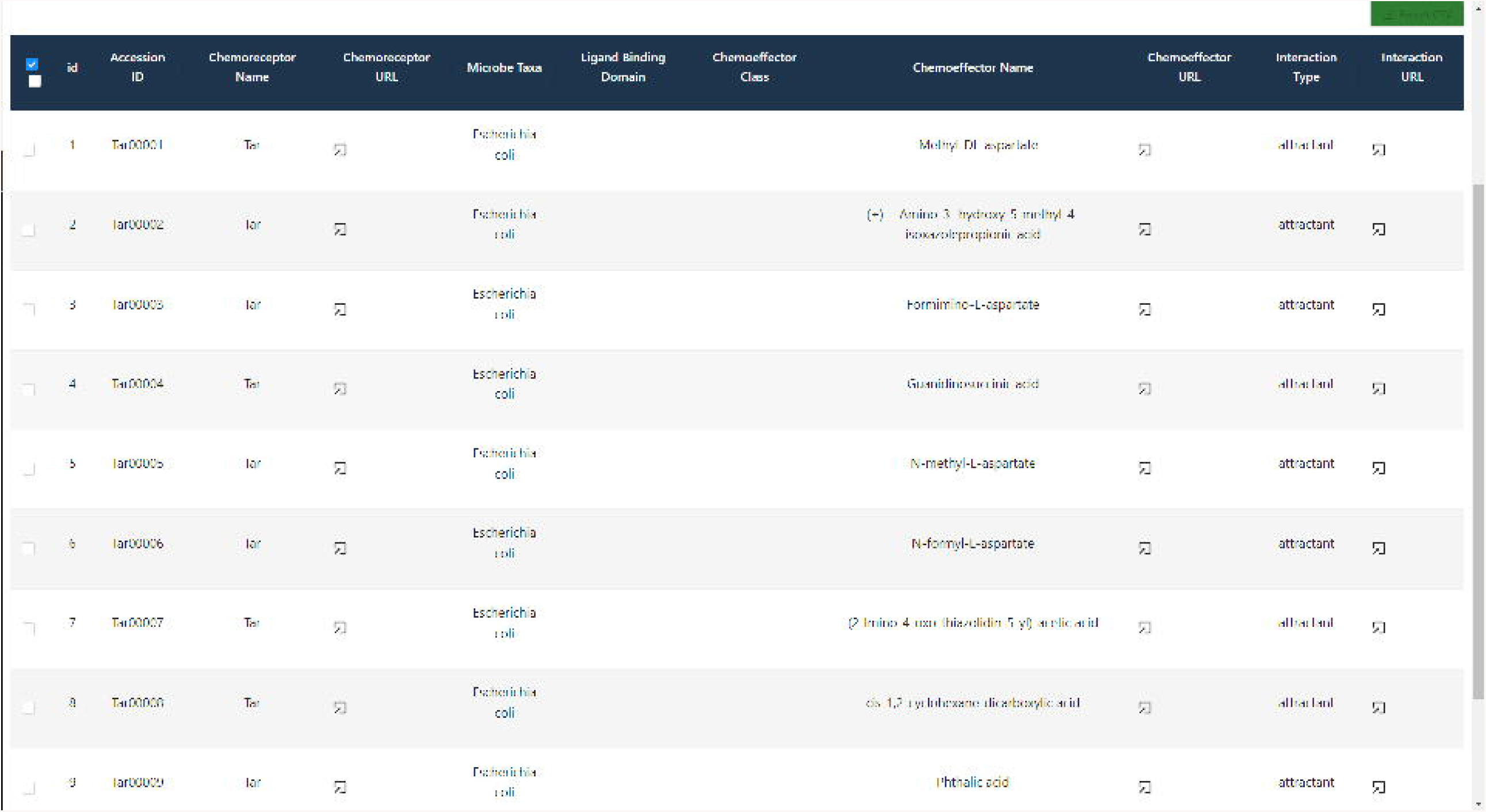

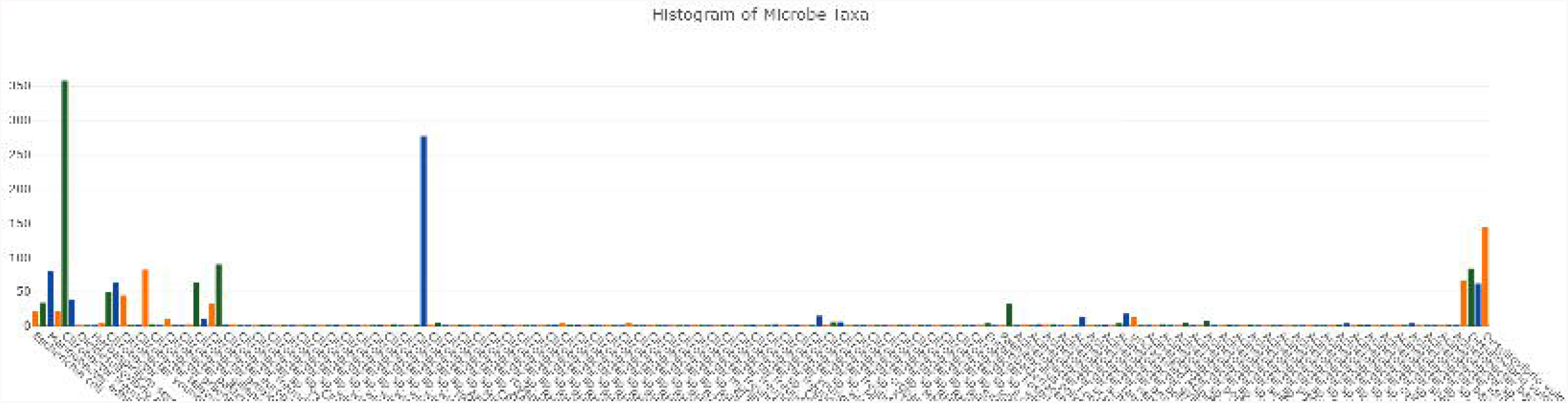

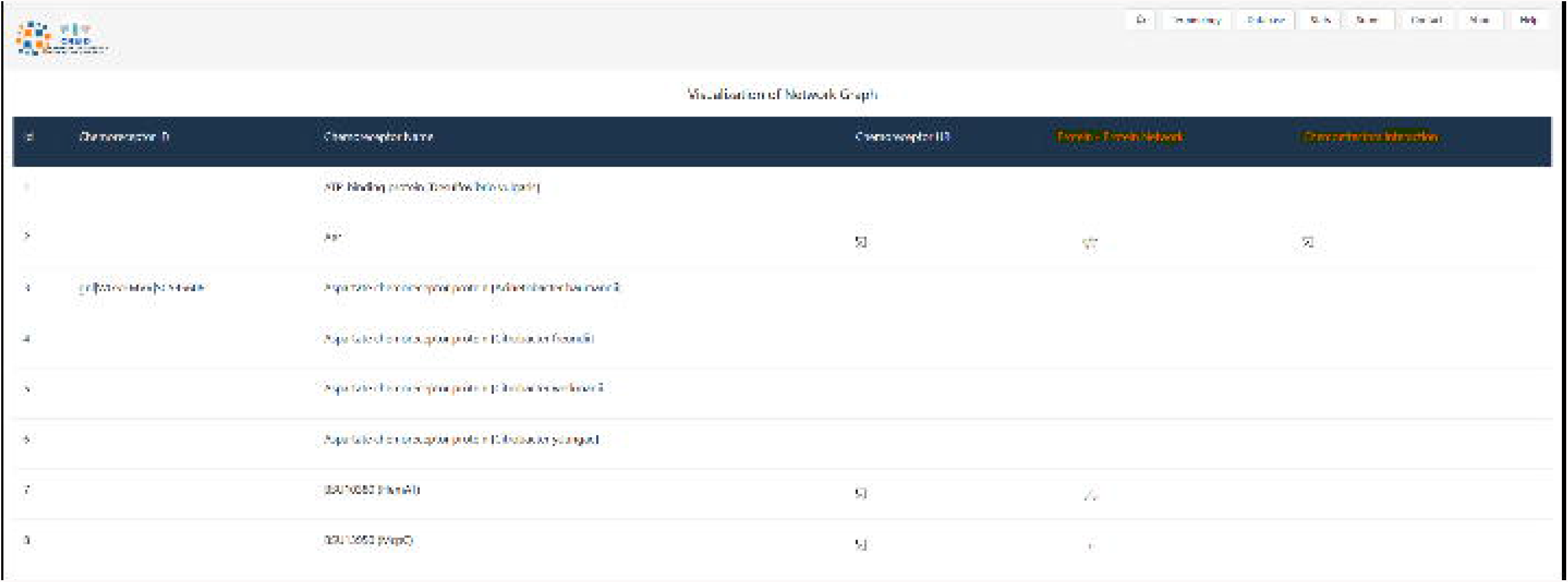

