## Supplementary figures and images for "CREID – A ChemoReceptor-Effector Interaction Database"

### Supplemental GIF

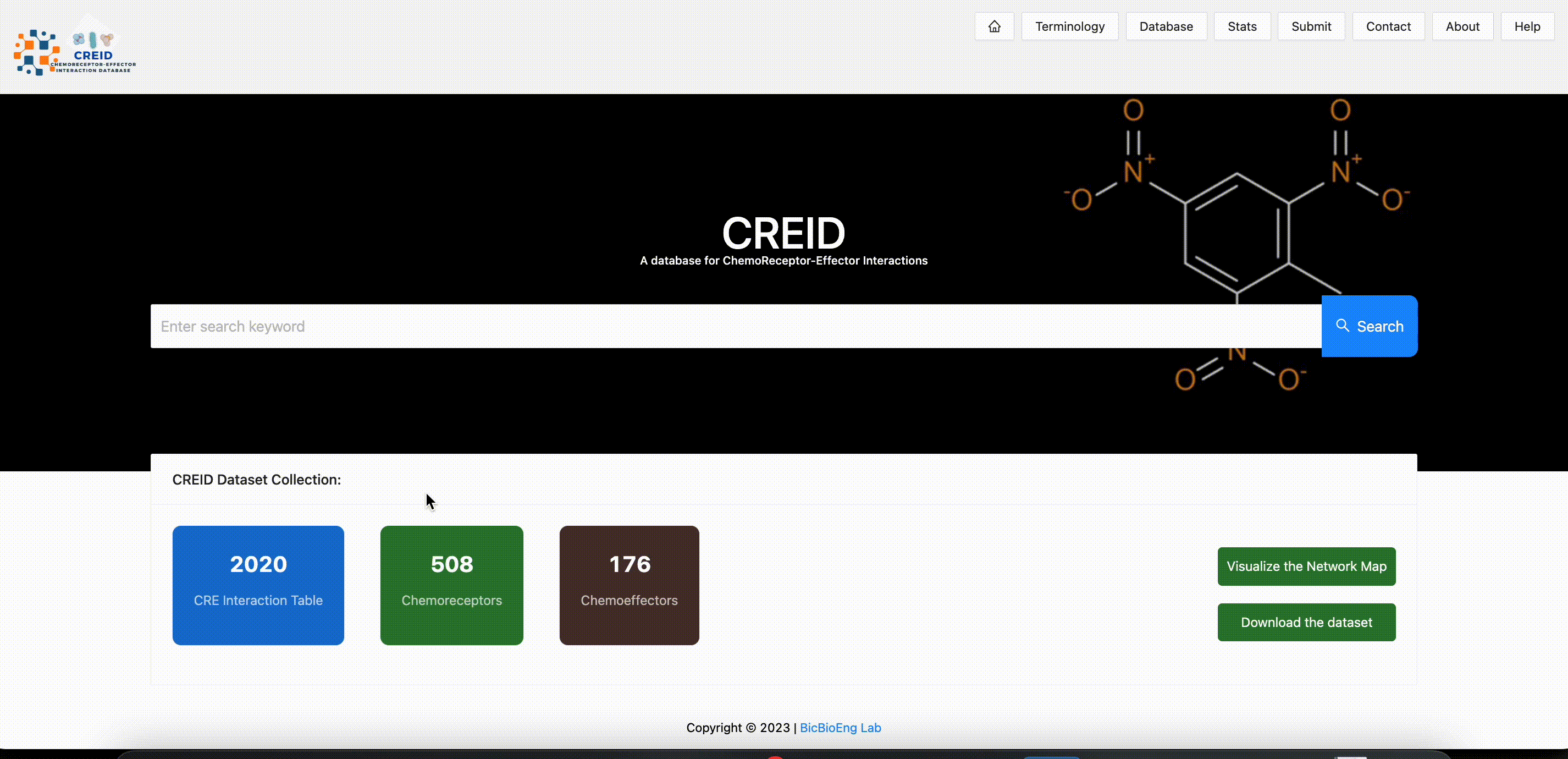
